# Mutation N501Y in RBD of Spike Protein Strengthens the Interaction between COVID-19 and its Receptor ACE2

**DOI:** 10.1101/2021.02.14.431117

**Authors:** Fang Tian, Bei Tong, Liang Sun, Shengchao Shi, Bin Zheng, Zibin Wang, Xianchi Dong, Peng Zheng

## Abstract

SARS-CoV-2 is spreading around the world for the past year. Enormous efforts have been taken to understand its mechanism of transmission. It is well established now that the receptor-binding domain (RBD) of the spike protein binds to the human angiotensin-converting enzyme 2 (ACE2) as its first step of entry. Being a single-stranded RNA virus, SARS-CoV-2 is evolving rapidly. Recently, several variants such as B.1.1.7, B.1.351, and P.1, with a key mutation N501Y on the RBD, appear to be more infectious to humans. To understand its mechanism, we combined cell surface binding assay, kinetics study, single-molecule technique, and computational method to investigate the interaction between these RBD (mutations) and ACE2. Remarkably, RBD with the N501Y mutation exhibited a considerably stronger interaction characterized from all these methodologies, while the other two mutations from B.1.351 contributed to a less effect. Fluorescence-activated cell scan (FACS) assays found that RBD N501Y mutations are of higher binding affinity to ACE2 than the wild type. Surface plasmon resonance further indicated that N501Y mutation had a faster association rate and slower dissociation rate. Consistent with the kinetics study, atomic force microscopy-based single-molecule force microscopy quantify their strength on living cells, showing a higher binding probability and unbinding force for the mutation. Finally, Steered Molecular Dynamics (SMD) simulations on the dissociation of RBD-ACE2 complexes revealed that the N501Y introduced additional π-π and π-cation interaction for the higher force/interaction. Taken together, we suggested that the reinforced interaction from N501Y mutation in RBD should play an essential role in the higher transmission of COVID-19 variants.

## Introduction

Over the past 20 years, coronavirus has posed severe threats to public health. In 2003, severe acute respiratory syndrome coronavirus (SARS-CoV-1) emerged in humans from an animal reservoir and infected over 8000 people with a ∼10% fatality rate^1, 2^. Middle East respiratory syndrome coronavirus (MERS-CoV) has infected over 1,700 people with a ∼36% fatality rate since 2012^3^. In late December 2019, a novel coronavirus, called severe acute respiratory syndrome coronavirus 2 (SARS-CoV-2), has been identified as the cause of an outbreak of a new respiratory illness named COVID-19. SARS-CoV-2 has caused more than two million deaths. Considerable effects have been taken to understand its molecule mechanism.

Coronaviruses are large, enveloped, positive-stranded RNA viruses belonging to the coronaviridae family, and they can be classified into four genera: alpha-coronavirus, beta-coronavirus, gamma-coronavirus, and delta-coronavirus^4^. SARS-CoV-2, SARS-CoV-1, and MERS-CoV are all beta-coronaviruses and only infect mammalians^5^. An envelope-anchored spike protein is capable of mediating coronavirus entry into host cells by first binding to a specific host receptor and then fusing viral and host membranes^5, 6^. The coronavirus spike protein, a class I fusion protein, is synthesized as a precursor single polypeptide chain consisting of three segments: a large ectodomain, a single-pass transmembrane anchor, and a short intracellular tail (Fig. 1a). After interacting with the host receptor, the spike protein is cleaved into an amino-terminal subunit (S1) and a carboxyl-terminal subunit (S2) by host furin-like proteases^6-8^. The receptor-binding domain (RBD) located in the C-terminal of the S1 subunit (S1 CTD) is responsible for recognizing and binding the host receptor and is critical in determining cell tropism, host range, and zoonotic transmission of coronaviruses^5, 7^. The S2 subunit contains a hydrophobic fusion loop and two heptad repeat regions (HR1 and HR2) and is in charge of membrane-fusion. The previous cryo-EM examination has illuminated the prefusion and postfusion structures of SARS-CoV-1 and SARS-CoV-2 spike protein, which reveal the molecular mechanism of coronavirus infection. The spike protein forms a clove-shaped homo-trimer with three S1 heads and a trimeric S2 stalk. Structural comparisons indicated that spike protein utilizes the CTD1 (N-terminal domain in CTD) as the receptor-binding domain, which changes the “down” conformation to the “up” conformation and then converts the inactivated state to the activated state to allow for receptor binding and possibly also initiate subsequent conformational changes of the S2 subunits to mediate membrane fusion^6-12^. The interaction of spike RBD of SARS-CoV-2 and ACE2 is the first step of viral entry into the host cell, and the great majority of vaccines and neutralizing antibodies are developing fast targeting for this region^13, 14^.

**Figure 1.**
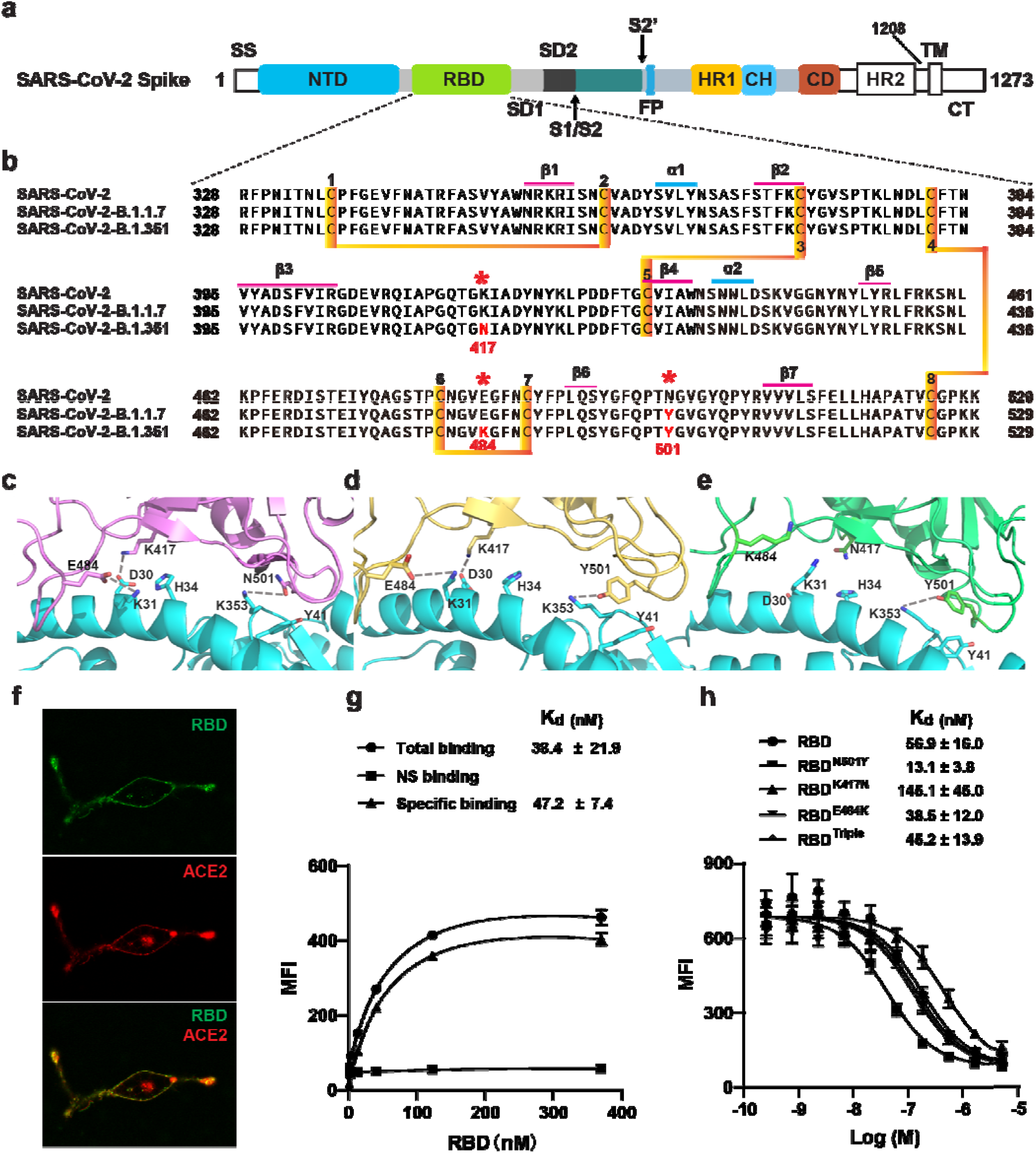
Two SARS-CoV-2 variants bound to ACE2 with higher affinity. a) Domain architecture of the SARS-CoV-2 spike monomer. NTD, N-terminal domain; SD1, subdomain 1; SD2, subdomain 2; FP, fusion peptide; HR1, heptad repeat 1; CH, central helix; CD, connector domain; HR2, heptad repeat 2; TM, transmembrane region; CT, C-terminal. b) Sequence alignment of RBD from SARS-CoV-2, B.1.1.7, and B.1.351 variants spike proteins. The N501Y, K417N, and E484K mutations are shown in red with a *. Cysteines forming disulfide bonds are marked in yellow. c-e) The interface of ACE2 (cyan) in complex with Spike RBD from SARS-CoV-2 (violet), B.1.1.7 lineage (orange), and B.1.351 line-age (green). Residues 501, 417, and 484 from RBD and RBD mutants, and contacting residues (D30, K31, H34, Y41, and K353) from ACE2 were showed in ball and sticks. Hydrogen bonds are shown in dash lines. f) Representative images of ACE2-mCherry (red) HEK293 cells stained with 100 nM AlexaFluor488-labeled RBD (green). g) Saturated binding of AlexaFluor488-labeled RBD to cell surface ACE2. NS, non-specific. h) Series-diluted RBD and RBD mutants were incubated with ACE2-expressing cells in the presence of AlexaFluor488-labeled RBD protein (100 nM). Concentrations used for unlabeled RBD and RBD mutants were from 5 μM to 0.25 nM with 3-fold dilution. K_d_ values were calculated using the Cheng-Prusoff equation.

Recently, several variants were found with increased transmissibility. The variant (B.1.1.7 lineage) was first detected in the United Kingdom (UK) in September 2020, and another variant (B.1.351 lineage)was first detected in October 2020 in the Republic of South Africa (RSA)^15-17^. They both carry an N501Y mutation in the RBD region, and the B.1.351 lineage has two more mutations (K417N, E484K) within the RBD region (Fig. 1b-c)^18, 19^. Recent data suggest that the FDA-authorized mRNA vaccines continued to induce a high level of neutralization against B.1.1.7 variant, but a lower level against B.1.351 variants. Several researchers assessed the neutralization potency of numerous antibodies against the two new variants. Their data suggest that some neutralizing antibodies in Phase II/III clinical trials were not able to retain their neutralizing capability against the B.1.351 variant. Since these mutations are within the RBD region, understanding the new variants’ binding mechanism to ACE2 receptor is of great value.

The structure of the RBD-ACE2 complex showed that extensive interactions have formed between them^20, 21^. To understand the potential role of these mutants binding to ACE2, we combine cell surface binding assay, kinetics study, single-molecule biophysical method, and SMD simulations to study the interaction of RBD mutants and ACE2 (Fig. 1d-e). Our results reveal the molecular mechanism of increased transmissibility of two SARS-CoV-2 variants by identifying the key mutation N501Y, which could be valuable for further developing vaccines and neutralizing antibodies against the SARS-CoV-2 virus mutation.

## Results

### Cell surface binding of RBD to ACE2

To elucidate the interaction between the RBD from SARS-CoV-2 variants and ACE2, we performed the cell surface-binding assay. ACE2 with a mCherry fused at the C-termini was transfected into HEK293 cells. Confocal microscopic images showed that ACE2 was located mainly in the cell membrane and endoplasmic reticulum (Fig. 1f). ACE2-mCherry positive cells were stained with AlexaFluor488 labeled-RBD, and the overlay images showed the co-localization of RBD and ACE2 on the cell surface (Fig. 1f). This visualized interaction laid the foundation for the performance of the following biochemical and biophysical experiments.

To measure the binding affinity of RBD from SARS-CoV-2 to ACE2 on the cell surface, saturation binding was performed using fluorescence flow cytometry by titration of Alexa488-RBD without washing the cells, which yields a K_d_ about 50 nM (Fig. 1g). Furthermore, we performed the competition-binding assay by titrating unlabeled RBD to compete with 100 nM Alexa488-RBD, which also showed a similar affinity of 50 nM (Fig. 1h).

### N501Y mutation slowed down the dissociation from the ACE2 receptor

To determine the role of the aforementioned mutations to the receptor, we first compared all the RBD mutants to wild-type RBD on the cell surface by competition binding assay (Fig. 1h). N501Y mutation from B.1.17 variant showed a 4-fold higher affinity than wild-type RBD on the cell surface. Mutation K417N or E484K showed a slightly weaker or similar affinity to cell surface ACE2, while triple mutation has a similar affinity as wild type. This demonstrated that N501Y mutation is the key residue to the higher affinity binding.

To further understand the change of kinetics of the mutations compared to wild-type RBD, we further performed surface plasmon resonance (SPR) on the immobilized RBD or RBD mutants with ACE2 as an analyte (Figure 2a-c, thin black lines). Compared to RBD, both RBD^N501Y^ and RBD^triple^ showed a 10-fold higher affinity contributed by a significantly lower off-rate and slightly higher on-rate (Fig. 2). Two other amino acid mutations (K417N and E484K) showed less impact on ACE2 binding, further verified by two single point mutants (Fig. S1). This result again emphasized the role of N501Y instead of the other two mutations for the higher binding affinity by slowing the dissociation rate from its receptor.

**Figure 2.**
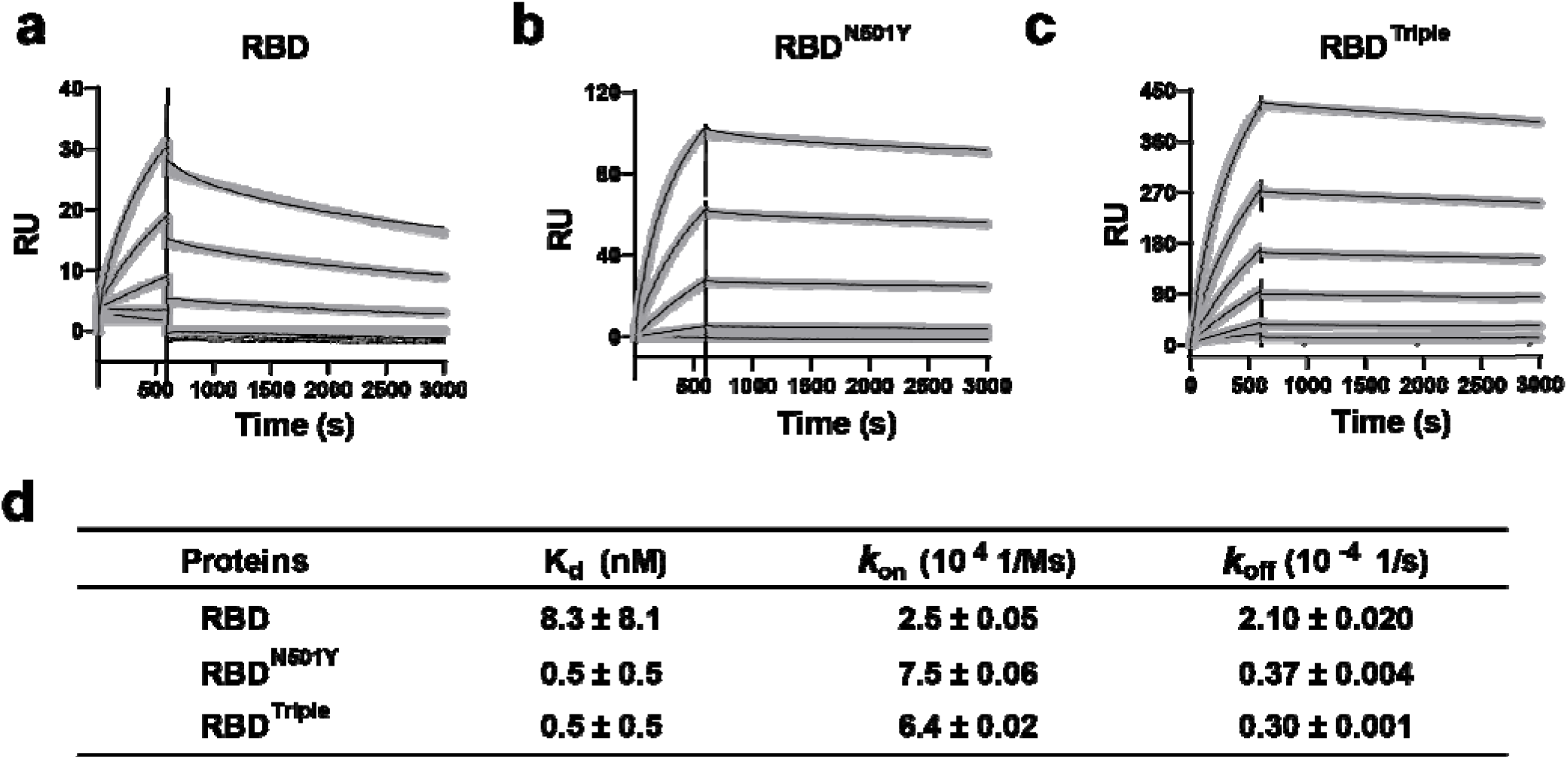
Kinetics of RBD and RBD mutants bound to ACE2 protein. a-c) Surface plasmon resonance (SPR) sensorgrams (thin black lines) were shown with fits (thick gray lines). Concentrations used for ACE2 protein were 50, 20, 10, 5, 2, and 1 nM respectively. Values were fitted to the 1:1 binding model. d) K_d_ and kinetic rates were showed as fit ± fitting error.

### AFM showed a higher binding probability and strength for the two mutants containing N501Y

In addition to classic ensemble measurements, we used atomic force microscopy-based single-molecule force spectroscopy (AFM-SMFS) to directly measure the binding strength between the three different RBDs and ACE2 on a living cell, respectively^22, 23^. As powerful single-molecule nanotechnology, AFM-SMFS can manipulate single molecule or molecules mechanically similar to optical and magnetic tweezers ^24-28^. It has been widely used to study the protein mechanics and protein-protein interaction, including the interaction between spike proteins of viruses and living cells^14, 29-33^. Previous AFM experiment has identified the binding events between the wild-type RBD and human ACE2 transfected on A549 cells, obtaining their binding probability and unbinding force/kinetic under a proper contact time and force^31^. In our work, a single RBD is site-specifically immobilized to a peptide-coated AFM tip via an enzymatic ligation (Fig. 3a, step (1), SI)^34-38^. An N-terminal GL sequence is present in the peptide. And a C-terminal NGL is added to the three RBDs. These two sequences can be recognized and ligated by protein ligase *Oa*AEP1 into a peptide bond linkage, and the RBD is attached to the tip for AFM measurement^34, 39^. Then, we used the ACE2-mCherry transfected HEK293 cells as the target cell, which is immobilized on a petri dish coated with poly-D-lysine. With the help of fluorescence, we targeted the transfected cell for measurement (Fig. 3b).

**Figure 3.**
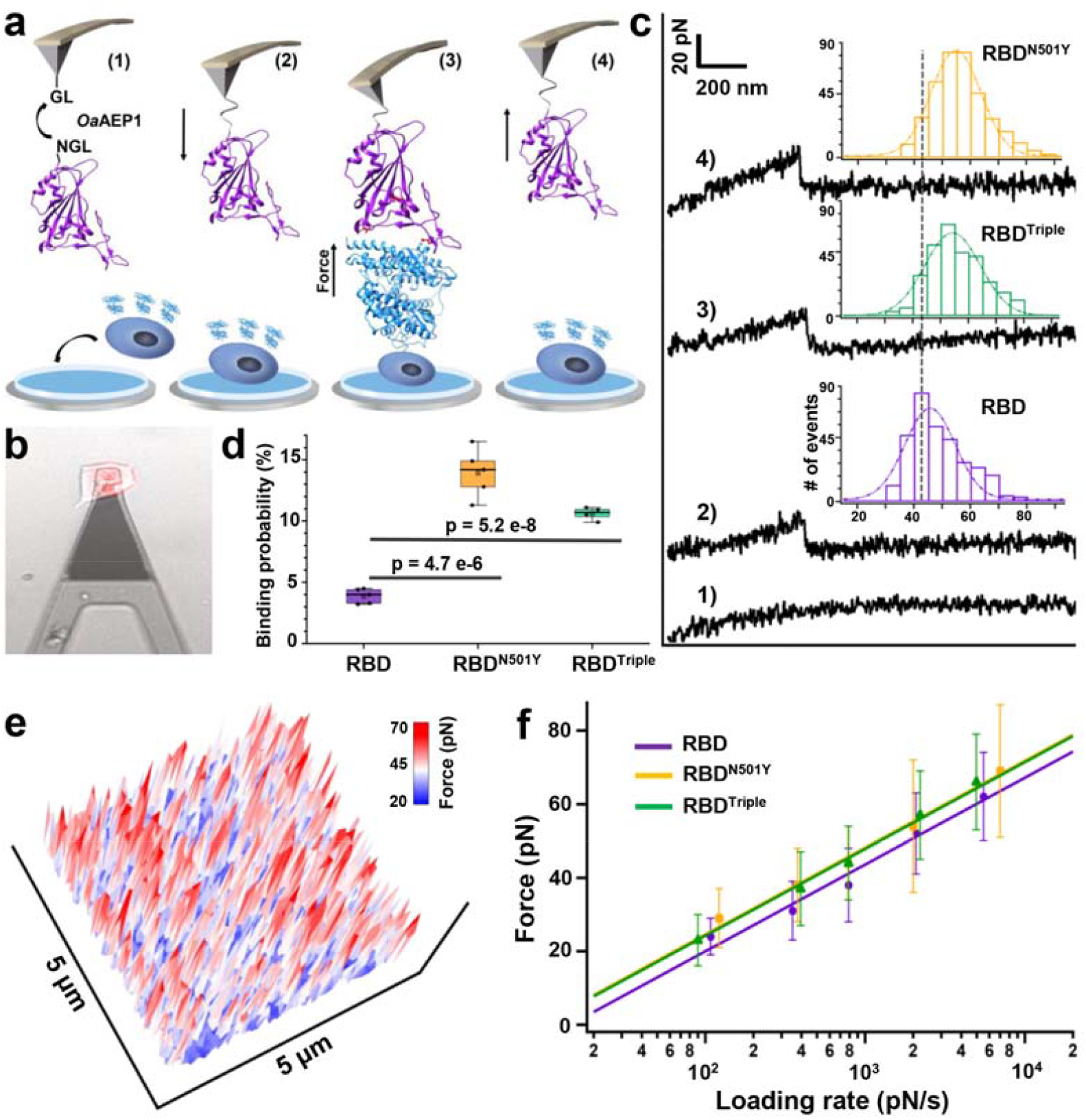
AFM-SMFS experiment to quantify the strength between RBDs and ACE2 on the living cell. a) Schematics of AFM-SMFS measurement processes show how the interaction is quantified. RBD with an N-terminal NGL is immobilized on a GL-coated AFM tip by ligase *Oa*AEP1, which recognizes the two sequences and ligates them into a peptide bond 1). By approaching the AFM tip to the target cell 2), RBD binds to ACE2 3). Then the tip retracts, and the complex dissociates finally, leading to an unbinding force peak 4). b) Image showed the reddish ACE2-mCherry transfected HEK293 cell being measured under the AFM tip by an inverted fluorescent microscope. c) Representative force-extension curves show no binding event (curve 1) and specific binding events between RBD-ACE2 complexes with an unbinding force peak (curves 2-4). From the force histogram (inset), RBD^N501Y^ and RBD^Triple^ showed higher forces as 57 pN and 56 pN than the RDB (49 pN). d) Box plot of the specific binding probabilities between the three RBDs and cell indicated a higher probability for the two mutants from AFM experiments under all five different velocities. The box indicates the 25^th^ and 75^th^ percentiles. e) 3D AFM force mapping of the cell surface showed the unbinding force distribution. f) The plot between loading rate and most probable unbinding forces from the complexes showed a linear relationship. The data is fitted to the Bell-Evans model to extract the off-rate.

By moving the AFM tip towards the cell, RBD contacts the cell and binds to the ACE2 on the surface (steps (2)-(3)). Then, the tip retracts at a constant velocity and pulls the complex apart by breaking all the interactions, leading to a force-extension curve with a force peak from the unbinding of the RBD-ACE2 complex (steps (3)-(4), Fig. 3c)^40^. If the RBD does not bind to the ACE2 receptor, a featureless curve will be observed (Fig. 3c, curve 1). Finally, the tip moves to another spot (65 nm away) on the cell and repeats the cycle for tens of hundreds of times, leading to a force map with unbinding force distribution of RBD for the cell surface (Fig. 3e)^41-43^.

For example, 2815 pieces of force-extension curves from probing the RBD^N501Y^-functionalized AFM tip to the ACE2-transfected cell have been obtained under the pulling speed of 5 µm/s. Based on its unbinding force (>20 pN), 14% of the events showed a specific interaction between RBD^N501Y^ and ACE2 (Fig. 3d, Fig. S2b, curve 4). The same experiments and analysis for RBD (3.3%) and RBD^Triple^ (11.2%) were performed. The interaction between RBD and normal HEK293 cell (1.7%) was also measured as a control (Fig. S3a). The unbinding force for RBD, RBD^N501Y^ and RBD^Triple^ is 49±11 pN (*n*=349), 57±18 pN (*n*=394) and 56±12 pN (*n*=312), respectively (Fig. 3b). Both RBD mutants showed a higher binding probability and unfolding force than the wild type, while the two variants’ properties are similar to each other (Fig. 3d&e, Fig. S3b-d).

Moreover, AFM-SMFS can obtain the unbinding kinetics. According to the Bell-Evans model, the externally applied force by AFM lowers the unbinding activation energy^44-47^. Thus, the binding strength of the ligand-receptor bond/interaction is proportional to the logarithm of the loading rate, which describes the effect of force applied on the bond over time. Thus, we pulled the RBD-ACE2 complexes at different velocities, and the relationship between unbinding forces and loading rate is plotted (Fig. 3f, Fig. S4). From the fit (SI), we can estimate the bond dissociation rate (*k*_off_) and the length scale of the energy barrier (Δ x_β_)^48^. Similarly, the *k*_off_ of the two RBD mutants is close to each other (0.030 s^-1^ and 0.035□s^-1^) but slower than the RBD (0.075□s^-1^).

### SMD simulations revealed a higher unbinding force and additional interactions for the complex with RBD mutant

There were no structural data available on the RBD mutants-ACE2 complexes, so we built models using the CHARMM36 force field^49^ (Fig. 1c, PDB: 6M0J). The obtained structures were further refined with MD simulations. Equilibration for 100 ns was performed using NAMD through its QwikMD interface^50,51^. Figure 1d,e shows the structure obtained after equilibration by MD simulations. Compared with the initial structure, the equilibrium results in a stable complex with no major changes in conformation.

To explore the possible molecular mechanism of the RBD-ACE2 complex dissociation under force, we performed SMD simulations to visualize the unbinding process during the AFM study (Fig. 4a, supplementary movies 1-3)^52-54^. The C-terminus of ACE2 was fixed, and the C-terminus of RBD was pulled at a constant velocity of 5.0 Å/ns (Fig. 4c-d, Fig. S5). In wild-type RBD-ACE2 interaction, N501 formed a hydrogen bond with K353 from ACE2. K417 and E484 from RBD formed salt bridges with D30 and K31 from ACE2, respectively (Fig. 4b). The RBD-ACE2 complex was ruptured under a force of ∼440 pN (Fig. 4a).

**Figure 4.**
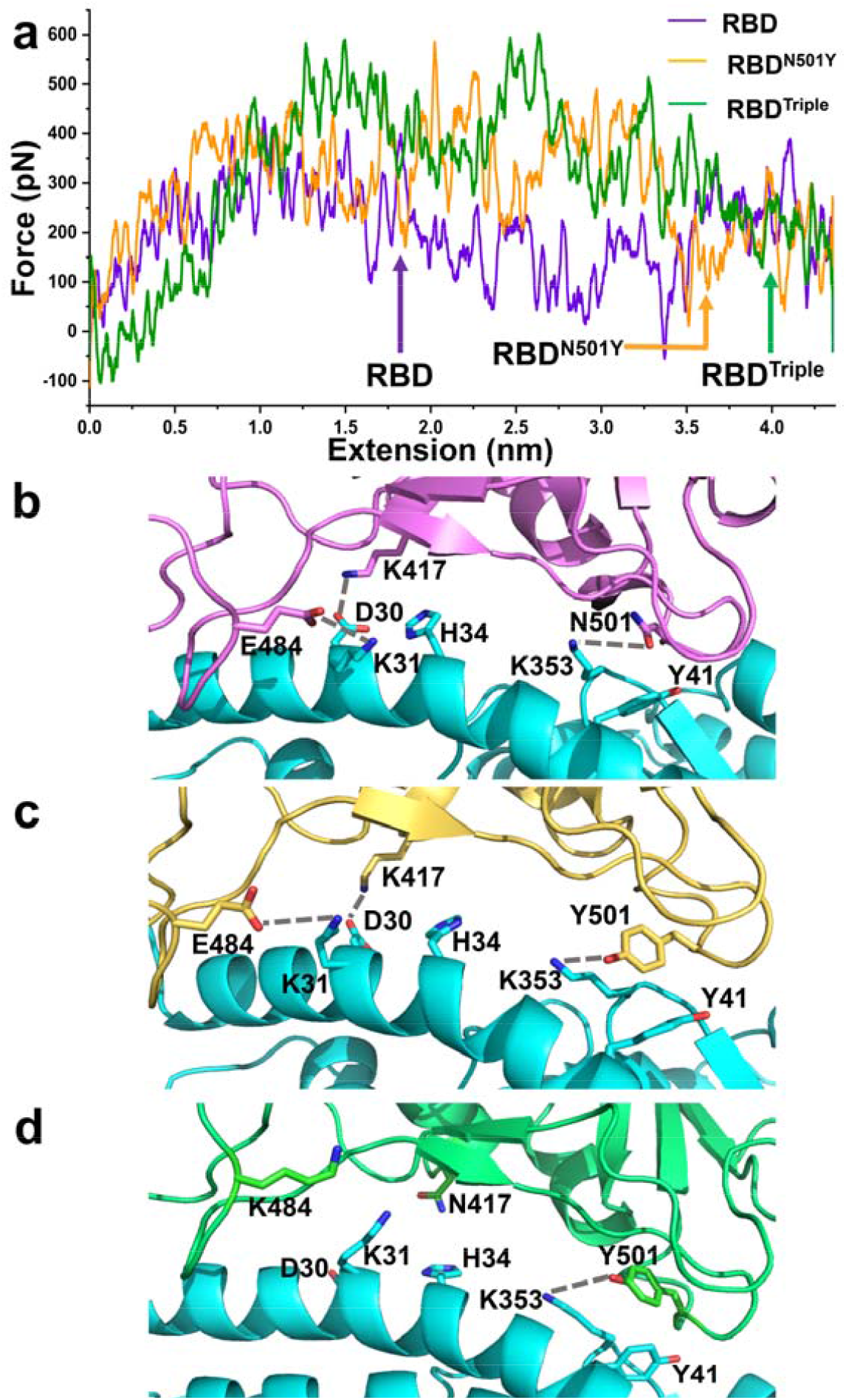
SMD simulations of the RBD-ACE2 complex dissociation. a) Force-extension traces of RBD-ACE2 (violet), RBD^N501Y^-ACE2 (orange), and RBD^Triple^-ACE2(green) complexes pulled at 5 Å/ns. Arrows indicate the key step during the dissociation of these complexes, of which structures are shown in b-d), respectively. ACE2 is colored in cyan. Key residues involved in the interaction between RBDs and ACE2 are depicted in the ball-and-stick model. Hydrogen bonds are indicated by dashed lines.

In the N501Y mutant, instead of the hydrogen bond with ACE2 K353, Y501 form a π-π interaction with Y41 and a π-cation interaction with K353 from ACE2, which could substantially reinforce the interaction. During the simulations, RBD^N501Y^ dissociated from ACE2 with a significant elevated unbinding force of ∼580 pN (Fig. 4a and 4c). Similarly, SMD simulations revealed that RBD^Triple^ might have a stronger contact than wild-type RBD with ACE2 too (Fig. 4a and 4d).

## Discussion

As coronaviruses are large, enveloped, positive-stranded RNA viruses, it has a remarkable mutation rate to evolve during the transmission. In this study, we combined the cell surface-binding assay, mechanical manipulation by AFM-SMFS, and molecular dynamics simulations to understand the behavior of key mutations recently detected in B.1.1.7 and B.1.351 variants for RBD binding. Among the three mutations, the N501Y mutation in RBD has the most significant role in binding and dissociation from the ACE2 receptor revealed by all the methodologies used in this work. Compared to the result of wild-type RBD, cell surface binding assay showed an increased binding affinity for RBD^N501Y^. SPR and AFM-SMFS measurements showed consistent results that RBD^N501Y^ had increased *k*_on_ and binding probability and decreased *k*_off,_ respectively. It is noted that SPR was performed between RBD and isolated ACE2 protein, while the other two measurements were performed between RBD and ACE2 on living cells. The more complex cell surface may account for the relatively weaker results from the cellular level, in which other proteins/receptors may interact with RBD in a non-specific and weaker way. Indeed, AFM unbinding results showed a fraction of non-specific interaction between RBDs and the un-transfected normal cell surface (∼5% for RBD monomer on HEK293 cell studied here, ∼10% for S1/RBD homotrimer on A549 cell)^17^. Indeed, previous AFM results on wild-type RBD showed a higher specific binding probability than us (∼20% vs. ∼15%), possibly due to different RBD construct and immobilization method and different A549 cell used^31, 34, 55^. Nevertheless, the specific unbinding force between RBD-ACE2 (∼50 pN) detected in the two experiments is similar. Moreover, the two mutants showed higher binding probability, higher unbinding force, and lower off-rate under the same condition in our studies.

The effect on the complex stability between RBD^N501^ and RBD^Triple^ was small, where two more mutations are present in RBD^Triple^. Indeed, FACS results showed that E484K contributed less to the interaction increment than N501Y, while K417N even decreased the interaction. Their effect may cancel due to the opposite effect from the two additional mutations, and N501Y is the dominant site. Thus, the combinations of all these three mutations in RBD^Triple^ lead to a similar effect as RBD^N501^. Indeed, a similar *k*_off_ value is obtained for RBD^N501^ and RBD^Triple^ from AFM-SMFS measurement.

It is noted that another SARS-COV-2 variant, P.1 lineage, is identified very recently in January 2021 in Brazil. Studies found that it may also affect the ability of antibodies to recognize and neutralize the virus as the two variants studied in this work. Interestingly, three key mutations (N501Y, E484K, and K417T) are found in its RBD, in which the N501Y mutation is still present. Consequently, we believe that the N501Y is a critical mutation to affect the transmission of COVID-19 by strengthening the interaction between RBD and ACE2. The strong interaction of N501Y mutation leads to the tighter binding of COVID-19 to the host cell to complete the membrane fusion or internalize the receptor together with the virus, leading the variant strain much more infective to humans.

Notably, nowadays, most vaccines are designed based on RBD or spike protein. Neutralizing antibodies from these vaccines are mainly targeting RBD to weaken its binding to ACE2. Here, we found that RBD with N501Y mutation has a 5-10 times higher affinity than wild-type RBD. Thus, a higher vaccine-induced antibody titer or neutralizing antibodies of higher affinity is needed to compete for the RBDN501Y-ACE2 interaction. This will make the current vaccine less effective than to other mutations, i.e., K417N or E484K.

## Materials and Methods

### Protein expression and purification

The genes were ordered from genscript Inc. The RBD construct contains the SAS-COV-2 spike protein (residues 319-591), followed by a GGGGS linker and an 8XHis tag in pcDNA3.4 modified vector (SI). Its mutants, including RBD^N501Y^, RBD^K417N^, RBD^E484K^, and RBD^Triple^ (N501Y, K417N, E484K), were generated using the QuikChange kit. Their sequences were all verified by direct DNA sequencing. A C-terminal NGL was added to the RBD, which is used for the AFM-SMFS experiment. The human ACE2 construct contains the ACE2 extracellular domain (residues 19-740) and an Fc region of IgG1 at the C-terminus.

All RDB and human ACE2 proteins were expressed in Expi293 cells with OPM-293 CD05 serum-free medium (OPM Biosciences). For protein purification of RBD with His-tag, culture supernatant was passed through a Ni-NTA affinity column (Qiagen), and Fc-tag ACE2 protein was purified using a protein affinity A column (Qiagen). Proteins were further purified by gel filtration (Superdex™ 200 Increase 10/30GL, GE Healthcare). The full-length ACE2 construct contains the ACE2 protein (residues 1-805), followed by a GGSGGGGS linker and a mCherry tag in pcDNA3.4 modified vector. ACE2 expression cell line was constructed by transient transfection of HEK293 cells and used for flow cytometry and AFM in this study.

### Confocal microscopy

A confocal microscope was used to detect the binding between SARS-CoV-2 RBD and the ACE2 receptor on the cell surface. Briefly, ACE2-mCherry cells were seeded in a poly-D-lysine precoated confocal dish and stained with AlexaFluor488-labeled-RBD (100 nM) for 30 min at room temperature. After three washes, the samples were imaged on a confocal microscope (Zeiss LSM 710), and the images were prepared using the ZEN software.

### Cell-surface binding by FACS

For saturation binding, wide-type RBD protein was labeled with AlexaFluor®488 NHS Ester (Yeasen) according to the manufacturer’s instructions. ACE2 expression cells (mCherry positive) and control HEK293 cells (mCherry negative) were resuspended in PBS buffer and incubated with AlexaFluor488 labeled-RBD at 4L for 1 h and subjected to flow cytometry without washing. MFI of AlexaFluor488 was reported for total binding (mCherry positive) and non-specific binding (mCherry negative). K_d_ value was saturated fitted.

For the competitive binding, ACE2 expression cells were resuspended in PBS buffer and incubated with 100 nM AlexaFluor488 labeled-RBD in the presence of competitors (0-5000 nM of unlabeled-RBD and RBD mutants). The mixture was allowed to equilibrate for 1 h before flow cytometry (ThermoFisher Attune NxT) without washing. The binding of RBD mutants was reported by changes in MFI of AlexaFluor488. The decrease of MFI value was directly proportional to the increase in the concentration of competitors. Competition curves were fitted using nonlinear regression with top and bottom value shared. The IC_50_ value was converted to an absolute inhibition constant K_d_ using the Cheng-Prusoff equation^56^.

### Surface plasma resonance

SPR studies were performed using Biacore™ T200 (GE Healthcare). Purified RBD and RBD mutants were amine-immobilized on the CM5 chips. Purified ACE2 protein was injected at 20 μL/min in 0.15 M NaCl, 20 mM HEPES, pH 7.4. The surface was regenerated with a pulse of 25 mM HCl at the end of each cycle to restore resonance units to baseline. Kinetics analysis was performed with SPR evaluation software version 4.0.1 (GE Healthcare).

### AFM-SMFS experiment

Atomic force microscopy (Nanowizard4, JPK) coupled to an inverted fluorescent microscope (Olympus IX73) was used to acquire correlative images and the force-extension curve. The AFM and the microscope were equipped with a cell-culture chamber allowing maintaining the temperature (37□±□1□°C). Fluorescence images were recorded using a water-immersion lens (×10, NA 0.3). The sample was scanned using 32×32 pixels per line (1024 lines) and a sample number of 10500. AFM images and force-extension curves were analyzed using JPK data process analysis software. Optical images were analyzed using ImageJ software.

The D tip of the MLCT-Bio-DC cantilever (Bruker) was used to probe the interaction between RBD and ACE2 on the cell. Its accurate spring constant was determined by a thermally-induced fluctuation method^57^. Peptide linker, C-ELP_20_-GL, was used to functionalize AFM tips as previously described^34^. Typically, the tip contacted the cell surface for 400 *m*s under an indentation force of 450 pN to ensure a site-specifically interaction between RBD and ACE2 on a cell while minimizing the non-specific interaction.

### SMD simulation for the dissociation of RBD-ACE2 complex

The RBD structure model in complex with receptor ACE2 was taken from the Protein Data Bank (PDB: 6M0J). The two RBD mutations, RBD^N501Y^, and RBD^Triple^ have no structure available and were created using the CHARMM36 force field^58^. Each system underwent a similar equilibration and minimization procedure. The molecular dynamics simulations were set up with the QwikMD plug-in in VMD^59^, and simulations were performed employing the NAMD molecular dynamics package^60^. The CHARMM36 force field was used in all simulations. Simulations were performed with explicit solvent using the CHARMM TIP3P^46^ water model in the NpT ensemble. The temperature was maintained at 300.00 K using Langevin dynamics. The pressure was maintained at 1 atmosphere using Nosé-Hoover Langevin piston^61^. A distance cut-off of 12.0 Å was applied to short-range, non-bonded interactions, and 10.0 Å for the smothering functions. Long-range electrostatic interactions were treated using the particle-mesh Ewald method^62^. The motion equations were integrated using the r-RESPA^60^ multiple time-step scheme to update the short-range interactions every one step and long-range electrostatic interactions every two steps. The time step of integration was chosen to be 2 fs for all simulations. Before the MD simulations, all the systems were submitted to an energy minimization protocol for 1,000 steps. An MD simulation with position restraints in the protein backbone atoms was performed for 1 ns, with temperature ramping from 0k to 300 K in the first 0.25 ns, which served to pre-equilibrate the system before the steered molecular dynamics simulations. The RBD mutations were subjected to 100 ns of equilibrium MD to ensure conformational stability. All structures shown are from post-equilibration MD simulations.

To characterize the interaction between RBDs and ACE2, the SMD simulations with constant velocity stretching employed a pulling speed of 5.0 Å/ns, and a harmonic constraint force of 7.0 kcal/mol/Å^2^ was performed for 10.0 ns. In this step, SMD was employed by harmonically restraining the position of the C-terminus of ACE2 and pulling on the C-terminus of RBD or mutations. Each system was run at least three times. The numerical calculations in this paper have been done on the computing facilities in the High-Performance Computing Center (HPCC) of Nanjing University.

## Abbreviation

(COVID-19): Coronavirus disease-19;
(SARS-CoV-2): Severe acute respiratory syndrome coronavirus-2;
(RBD): Receptor-binding domain
(MERS-CoV): Middle East respiratory syndrome coronavirus;
(ACE2): Angiotensin-converting enzyme 2;
(SPR): Surface Plasmon Resonance;
(FACS): Fluorescence-activated cell Scan;
(AFM): Atomic Force Microscopy;
(SMFS): Single-molecule force spectroscopy;
(SMD): Steered Molecular Dynamics.

## Supporting Information

Supplementary materials, notes, figures, table, and movies are described in detail in the supporting information.

## Corresponding authors

, and

## Acknowledgment

This work was supported by the National Key Research and Development Program of China (Grant No. 2020YFA0509003, 2020YFA0710800), by Natural Science Foundation of Jiangsu Province (Grant No. BK20200058, BK20202004, BK20190275), by the National Natural Science Foundation of China (Grant No. 21771103, 21977047), by the Fundamental Research Funds for the Central Universities (Grant No. 14380205).

## Table of Contents Graphic

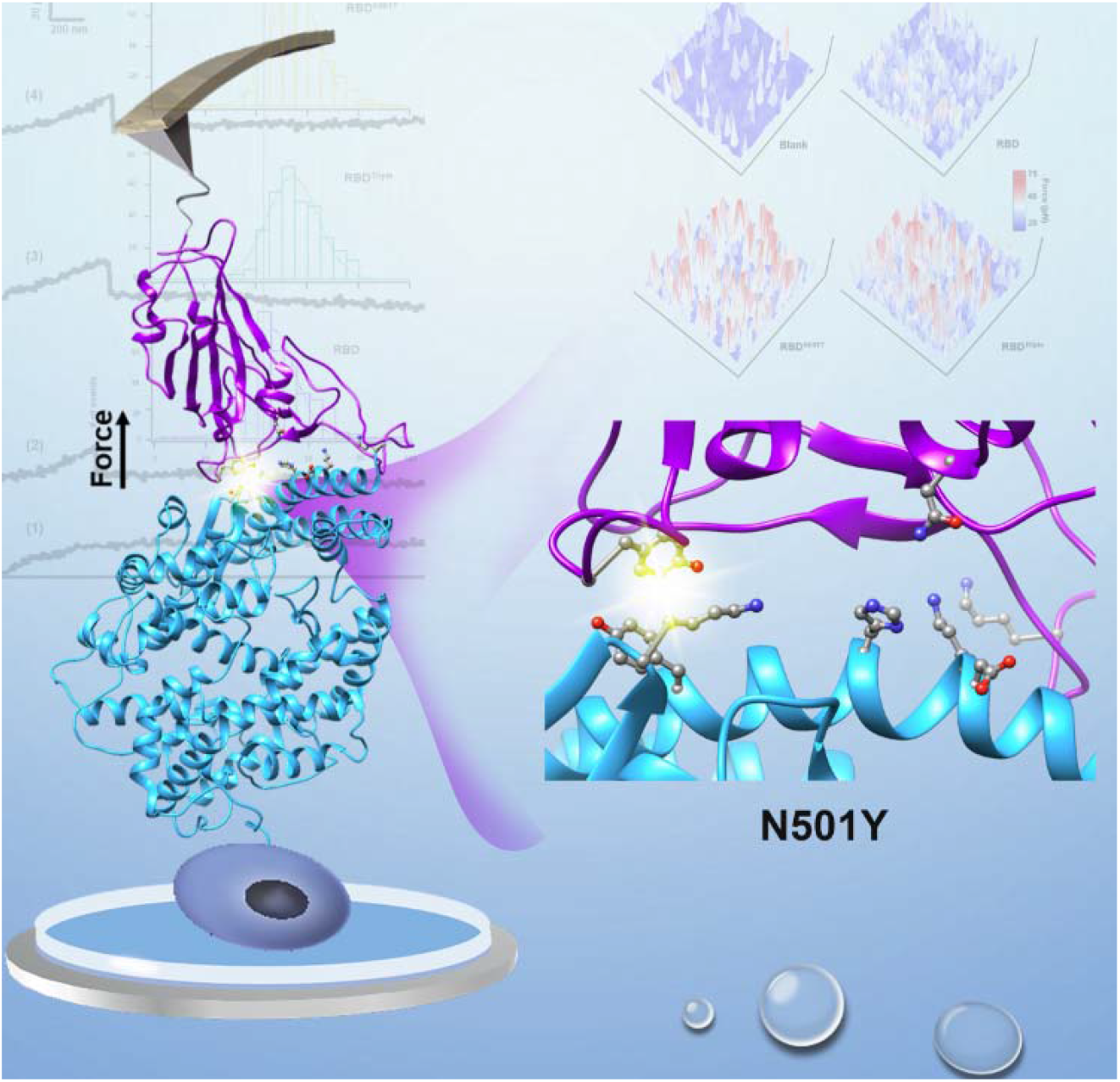

