## supplementary information for "Mutation N501Y in RBD of Spike Protein Strengthens the Interaction between COVID-19 and its Receptor ACE2"

This Supplementary Information Includes:

**Supplementary Methods	S3-S4**

Supplementary materials	S3

Protein engineering	S3

AFM tip functionalization	S3

AFM-SMFS experiment	S4

Bell-Evans model to extract kinetics	S4

**Supplementary Notes	S6**

Protein sequences	S6

**Supplementary Figures	S7-S9**

Figure S1.	S7

Figure S2.	S7

Figure S3.	S8

Figure S4.	S8

Figure S5.	S9

**Supplementary Table	S10**

**Supplementary Movies	S11**

**References	S12**

**Supplementary Methods**

**Supplementary materials.** All chemical reagents were purchased from commercial suppliers and used accordingly. (3-Aminopropyl) tiethoxysilane (APTES, Sigma Aldrich), sulfosuccinimidyl 4-(N-maleimidomehthyl) cyclohexane-1-carboxylate) (Sulfo-SMCC, Thermo Scientific) were stored and used in the dark. Aqueous solutions were prepared in Milli-Q water (18.2 MΩ/cm, 0.22-µm filter). For the culture of *E. coli*, Luria-Bertani (LB) medium and agar plates (Sangon Biotech) were used. Protein concentrations were routinely determined by Nanodrop 2000.

**Protein engineering.** *Oa*AEP1(C247A) is cysteine 247 to alanine mutant of asparaginyl endoproteases 1 from *oldenlandia affinis*, abbreviated as *Oa*AEP1^1^. ELP is the elastin-like polypeptides^2^. Protein *Oa*AEP1 and ELP were overexpressed in BL21(DE3) *E. coli* cells. The details for their expression and purification can be found in literatures^1-3^.

**AFM tip functionalization.** First, the maleimide group for cysteine coupling was added on the amino-functionalized AFM tip using the hetero-bifunctional crosslinker sulfo-SMCC from the reaction between amino and -NHS. Then, the peptide GL-ELP_20_-C was reacted to the maleimide. The long ELP serves as a spacer to avoid non-specific interaction between the tip and the cell surface. Finally, target RBDs with the C-terminal NGL sequence were site-specifically linked to the tip by ligase *Oa*AEP1, which recognized the N-terminal GL on the tip and the C-terminal NGL, forming a peptide bond^3^.

Specifically, the silicon nitride AFM cantilever (MLCT-BIO-DC, Bruker Corp.) was used. The tip was coated with the amino group by amino-silanization^4^. Briefly, the cantilevers were immersed in 1.5% (v/v) APTES toluene solution for 1 hour at room temperature in the dark and rinsed with ethyl alcohol. After drying, they were baked at 80 °C for 15 min and then cool down to RT. 200 μL of sulfo-SMCC (1 mg mL^−1^) in dimethyl sulfoxide (DMSO) was added and incubated for 2 h, protected from light. The cantilevers were washed with absolute ethyl alcohol to remove residual sulfo-SMCC. Finally, each cantilever was reacted with more than 50 μL of GL-ELP_20_-C and washed with 50 mL high-salt buffer (100 mM Tris, 1 M NaCl, pH 7.4), and dried. Consequently, these cantilevers are ready for RBD immobilization.

To add RBD on the AFM tip, the GL-functionalized cantilevers were incubated with 50 μL mixed solution of 60 μM RBD-NGL and 1 μM *Oa*AEP1 in the measurement buffer. The *Oa*AEP1-catalyzed coupling was performed in the measurement buffer (100 mM Tris, 100 mM NaCl, pH 7.4) at RT for ~30 minutes, forming a covalent NGL linkage between the AFM tip and RBD.

**AFM-SMFS experiment.** Nanowizard4 atomic force microscopy (JPK) is used to perform the AFM-SMFS unbinding experiments on cells. After calibration, the tip was pushed into the cell surface with a typical indentation force of 450 pN and a dwell time of 400 ms, and the specific interaction between RBD and ACE2 will be formed^5,6^. The pulling distance is 6 µm, and the sample number is 10500. Then, moving the tip up vertically at a constant velocity (5 µm/s, if not specified), the complex ruptured. Then, the tip moved to another place to repeat this cycle several thousands of times. As a result, a force-extension curve, possibly including the complex unbinding event, was obtained. The data was analyzed using data processing software (JPK).

**Bell-Evans model to extract kinetics.**

The RBD-ACE2 complex dissociation in the AFM experiment is a non-equilibrium process that can be modeled as an all-or-none two-state process with force-dependent rate constant *k*(F). The rate constant can be described by Bell-Evans’ model^7,8^:

$k\left( F \right)=k_{\mathrm{off}}exp(\frac{F\Delta x_{\beta}}{k_{b}T})$	(1)

*k*(*F*) is the complex dissociation rate constant under a stretching force of *F*, *k*_off_ is the dissociation rate constant under zero force, Δx_β_ is the distance between the bonded state and the transition state of dissociation.

For the dynamic force spectroscopy measurements, the slope $a$ of the force−extension curves immediately (2~3 nm) before the rupture event was first determined to obtain the average loading rate ($r=av$, where $v$ is the velocity in the rupture event). All the data were fitted with the Bell-Evans model (1), thus yielding the spontaneous rupture rate, and the distance from the bound state to the transition state with the following equation:

$F=\frac{k_{b}T}{\Delta x_{\beta}}\ln\left( \frac{\Delta x_{\beta}}{k_{\mathrm{off}}k_{b}T} \right)+\frac{k_{b}T}{\Delta x_{\beta}} ln(r)$	(2)

By performing the AFM unbinding experiment under five different pulling speeds, 0.5 µm/s, 1 µm/s, 3 µm/s, 5 µm/s, and 10 µm/s, the relationship between most probable rupture force and loading rate can be obtained on a log scale, which is fitted by a linear line as equation (2). Thus, the slope of this line can be used to calculate the Δx_β_, which is the distance between the bonded state and the transitional unbonded state, and the y-intercept is used to calculate the *k*_off._

**Supplementary Notes**

The protein sequence of **GL**-**(ELP)_20_**-**C**:

M**GL**HHHHHHGS**VPGEGVPGVGVPGVGVPGVGVPGVGVPGAGVPGAGVPGGGVPGGGVPGEGVPGEGVPGVGVPGVGVPGVGVPGVGVPGAGVPGAGVPGGGVPGGGVPGEG**RS**C**

The protein sequence of **RBD (or RBD mutants)-NGL**:

**RVQPTESIVRFPNITNLCPFGEVFNATRFASVYAWNRKRISNCVADYSVLYNSASFSTFKCYGVSPTKLNDLCFTNVYADSFVIRGDEVRQIAPGQTGK(N)IADYNYKLPDDFTGCVIAWNSNNLDSKVGGNYNYLYRLFRKSNLKPFERDISTEIYQAGSTPCNGVE(K)GFNCYFPLQSYGFQPTN(Y)GVGYQPYRVVVLSFELLHAPATVCGPKKSTNLVKNKCVNFNFNGLTGTGVLTESNKKFLPFQQFGRDIADTTDAVRDPQTLEILDITPCS**GGGGSHHHHHHHHGGS**NGL**

The protein sequence of **ACE2-***mCherry*：

**QSTIEEQAKTFLDKFNHEAEDLFYQSSLASWNYNTNITEENVQNMNNAGDKWSAFLKEQSTLAQMYPLQEIQNLTVKLQLQALQQNGSSVLSEDKSKRLNTILNTMSTIYSTGKVCNPDNPQECLLLEPGLNEIMANSLDYNERLWAWESWRSEVGKQLRPLYEEYVVLKNEMARANHYEDYGDYWRGDYEVNGVDGYDYSRGQLIEDVEHTFEEIKPLYEHLHAYVRAKLMNAYPSYISPIGCLPAHLLGDMWGRFWTNLYSLTVPFGQKPNIDVTDAMVDQAWDAQRIFKEAEKFFVSVGLPNMTQGFWENSMLTDPGNVQKAVCHPTAWDLGKGDFRILMCTKVTMDDFLTAHHEMGHIQYDMAYAAQPFLLRNGANEGFHEAVGEIMSLSAATPKHLKSIGLLSPDFQEDNETEINFLLKQALTIVGTLPFTYMLEKWRWMVFKGEIPKDQWMKKWWEMKREIVGVVEPVPHDETYCDPASLFHVSNDYSFIRYYTRTLYQFQFQEALCQAAKHEGPLHKCDISNSTEAGQKLFNMLRLGKSEPWTLALENVVGAKNMNVRPLLNYFEPLFTWLKDQNKNSFVGWSTDWSPYADQSIKVRISLKSALGDKAYEWNDNEMYLFRSSVAYAMRQYFLKVKNQMILFGEEDVRVANLKPRISFNFFVTAPKNVSDIIPRTEVEKAIRMSRSRINDAFRLNDNSLEFLGIQPTLGPPNQPPVSIWLIVFGVVMGVIVVGIVILIFTGIRDRKKKNKARSGENPYASIDISKGENNPGFQNTDDVQTSFDYKDDDDK-**

GGSGGGGS-M*VSKGEEDNMAIIKEFMRFKVHMEGSVNGHEFEIEGEGEGRPYEGTQTAKLKVTKGGPLPFAWDILSPQFMYGSKAYVKHPADIPDYLKLSFPEGFKWERVMNFEDGGVVTVTQDSSLQDGEFIYKVKLRGTNFPSDGPVMQKKTMGWEASSERMYPEDGALKGEIKQRLKLKDGGHYDAEVKTTYKAKKPVQLPGAYNVNIKLDITSHNEDYTIVEQYERAEGRHSTGGMDELYK*

**Supplementary Figures**

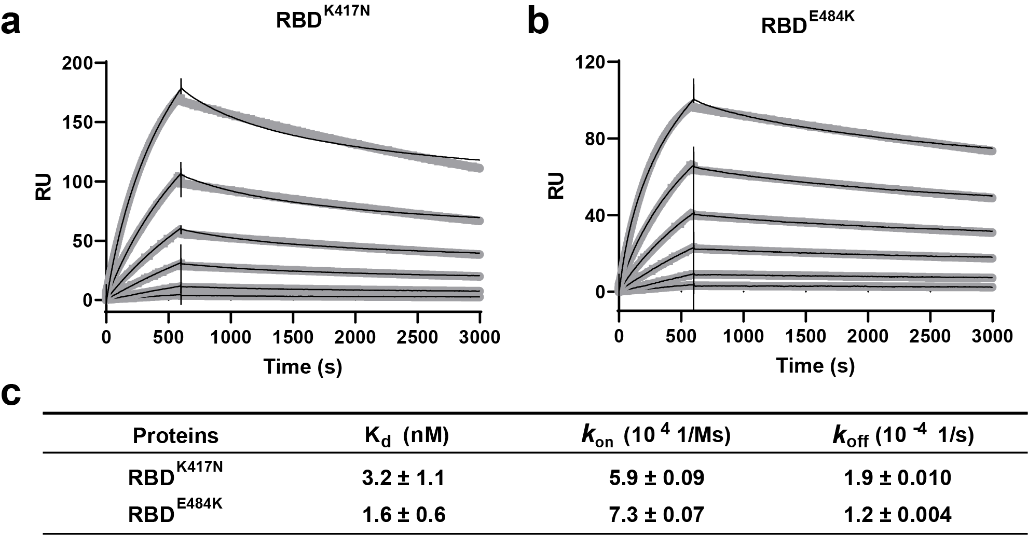

Figure S1. Kinetics of two RBD mutants bound to ACE2 protein. (a-b) SPR sensorgrams (thin black lines) were shown with fits (thick gray lines). Concentrations used for ACE2 protein were 50, 20, 10, 5, 2, and 1 nM respectively. Values were fitted to the 1:1 binding model. (d) K_d_ and kinetic rates were showed as fit ± fitting error.

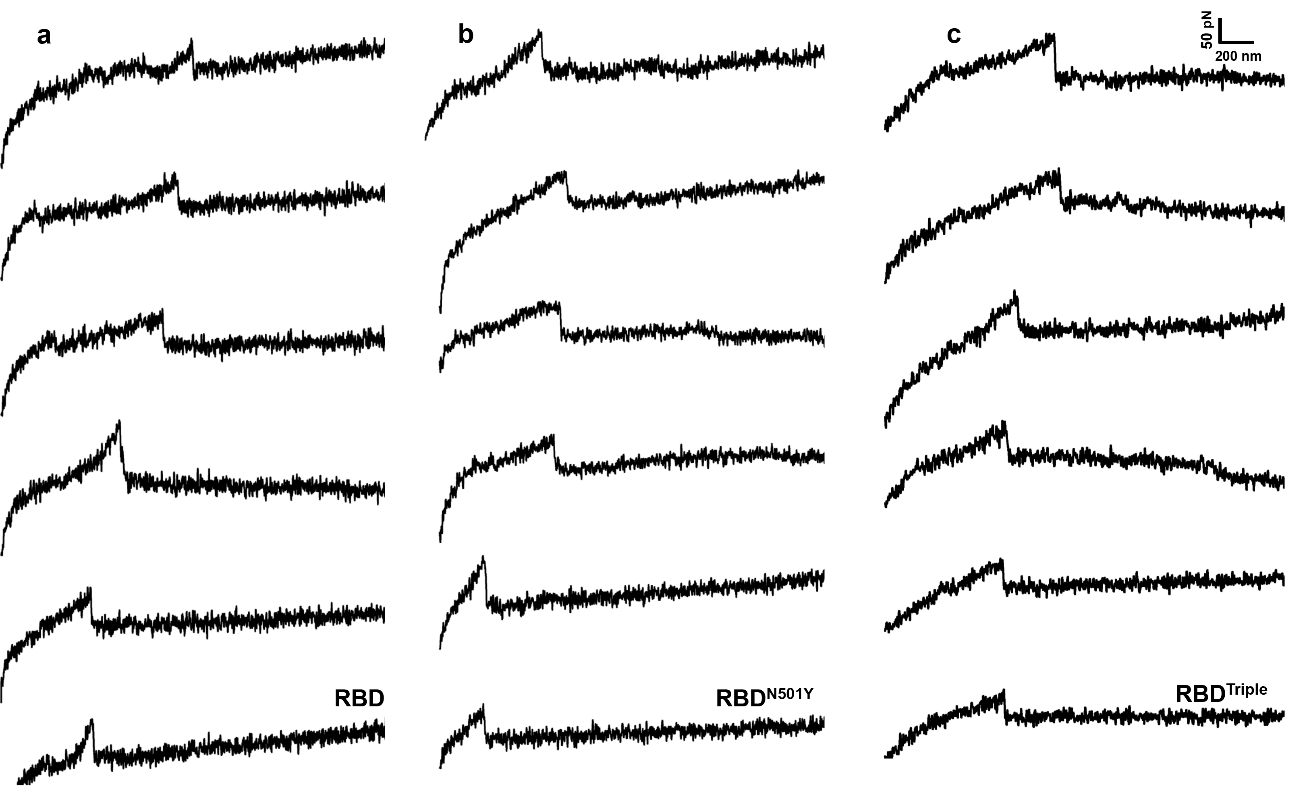

Figure S2. More representative unbinding force-extension curves for the three different RBD-ACE2 complexes a), RBD^N501^ b), and RBD^Triple^ c) are shown, respectively. A clear unbinding peak with force higher than 20 pN is present.

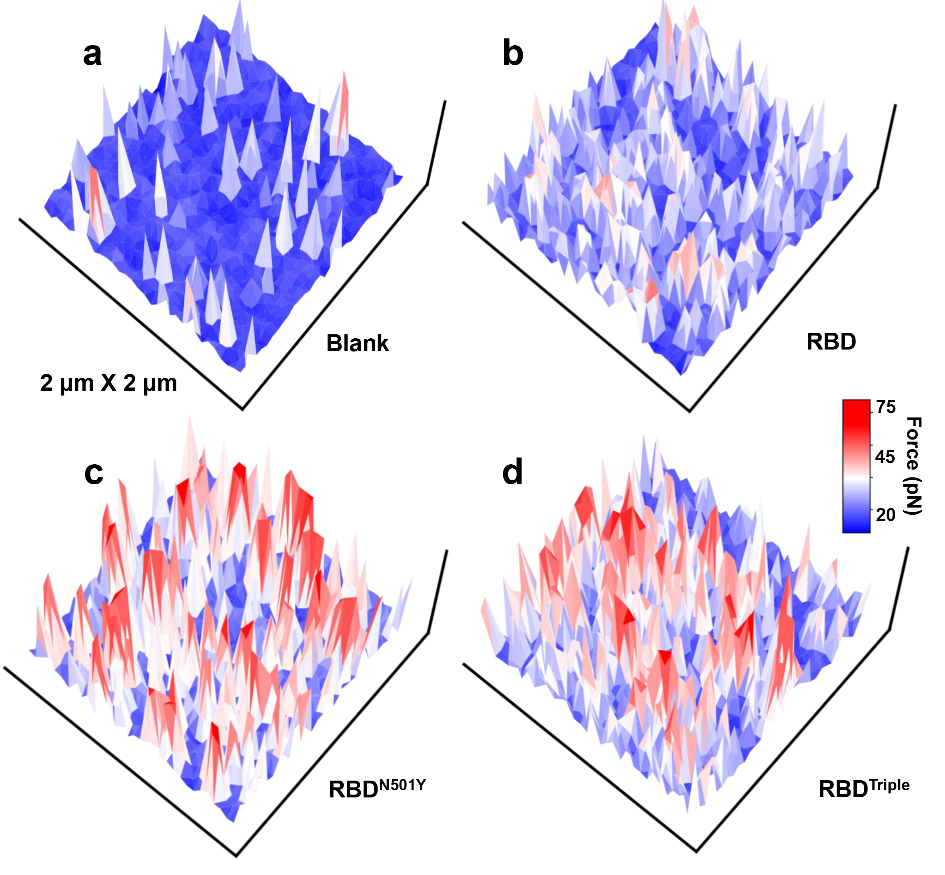

Figure S3. The force mapping results for the three complexes in a 2х2 µm area, blank on un-transfected HEK293 cell a), RBD b), RBD^N501^ c), and RBD^Triple^ d) are shown, respectively.

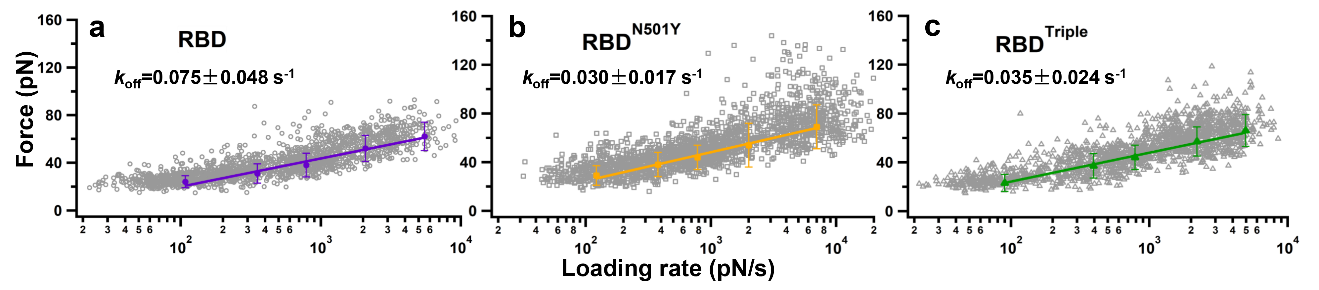

Figure S4. The plot between the raw unbinding force and loading rate of the three complexes RBD a), RBD^N501^ b), and RBD^Triple^ c) are shown, respectively. The gray marker is the force for an individual unbinding event under a specific loading rate. The five solid markers in each graph are the most probable unbinding force under the five different loading rates, respectively. The data is fitted by the Bell-Evans model to obtain the *k*_off_*_._*

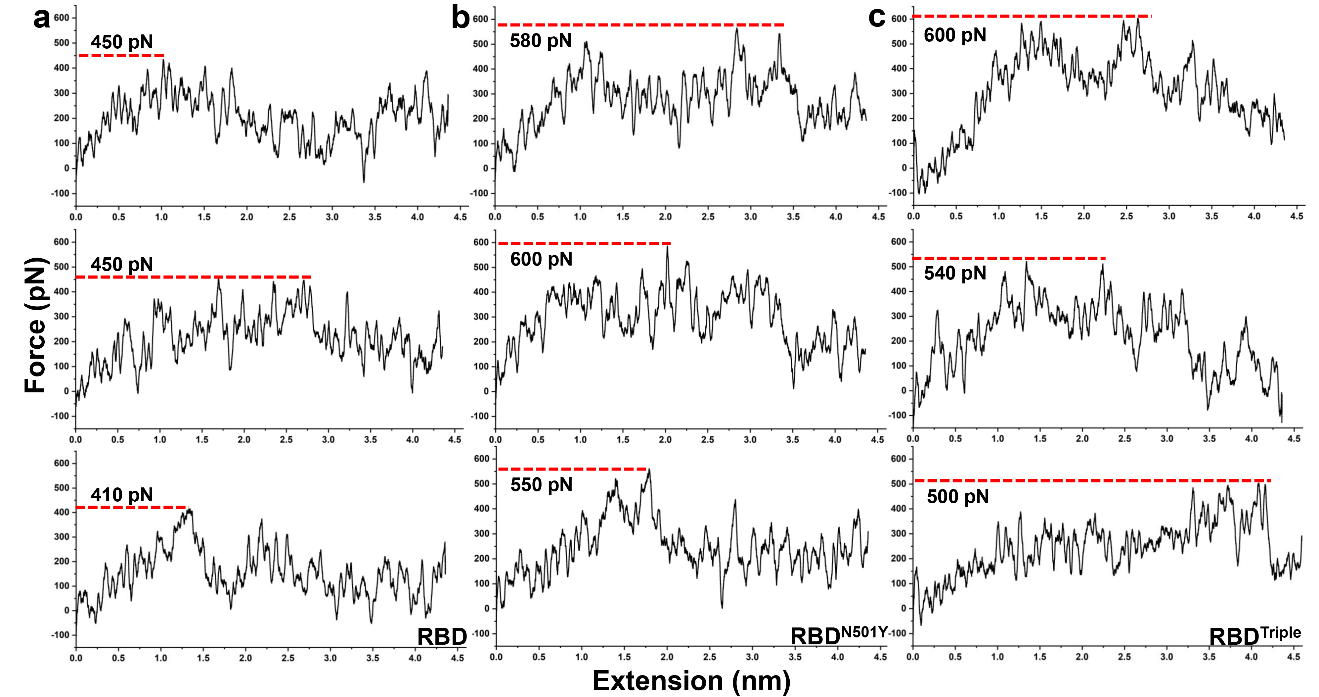

Figure S5. Three times SMD results of the different RBD-ACE2 complexes dissociation, RBD a), RBD^N501Y^ b), and RBD^Triple^ c) are shown, respectively.

**Supplementary Table**

Table S1. Summary of the spring constant (*k*) of all fifteen cantilevers

| **Velocity (µm/s)** | **RBD**  **(pN·nm^-1^)** | **RBD^N501Y^**  **(pN·nm^-1^)** | **RBD^Triple^**  **(pN·nm^-1^)** |
| --- | --- | --- | --- |
| **0.5** | **33.9/30.3/34.2** | **32.4/31.5** | **35.4/32.8** |
| **1** | **36.3/34.2/31.9** | **35.6/31.5** | **35.4/33.0** |
| **3** | **30.3/36.3/33.9** | **35.6** | **35.4** |
| **5** | **30.3/33.9** | **37.5/31.8/31.5** | **35.4/32.8/31.9** |
| **10** | **30.3/31.9/29.4** | **32.4/31.8/37.5** | **31.9/35.4/33.0** |

**Supplementary Movies**

**Movie S1.**

Representative SMD simulation of the RBD-ACE2 complex at a constant velocity of 5.0 Å/ns. RBD is colored in violet, ACE2 is colored in cyan, respectively. Residues 501, 417, and 484 from RBD, and contacting residues (D30, K31, H34, Y41, and K353) from ACE2 were showed in ball and sticks.

**Movie S2.**

Representative SMD simulation of the RBD^N501Y^-ACE2 complex at a constant velocity of 5.0 Å/ns. RBD^N501Y^ is colored in orange, ACE2 is colored in cyan, respectively.

**Movie S3.**

Representative SMD simulation of the RBD^Triple^-ACE2 complex at a constant velocity of 5.0 Å/ns. RBD^Triple^ is colored in green, ACE2 is colored in cyan, respectively.

**References：**
